# dGAMLSS: An exact, distributed algorithm to fit Generalized Additive Models for Location, Scale, and Shape for privacy-preserving population reference charts

**DOI:** 10.1101/2024.12.20.629834

**Authors:** Fengling Hu, Jiayi Tong, Margaret Gardner, Lifespan Brain Chart Consortium, Andrew A. Chen, Richard A.I. Bethlehem, Jakob Seidlitz, Hongzhe Li, Aaron Alexander-Bloch, Yong Chen, Russell T. Shinohara

**Author notes:** Correspondence: Fengling Hu <; 423 Guardian Dr, Philadelphia, PA 19104>. **CRediT author statement: Fengling Hu:** Conceptualization, Methodology, Software, Validation, Formal Analysis, Investigation, Data Curation, Writing – Original Draft, Writing – Review & Editing, Visualization. **Jiayi Tong:** Methodology, Validation, Formal Analysis, Resources, Investigation, Writing – Review & Editing. **Margaret Gardner:** Validation, Resources, Writing – Review & Editing. **Andrew A. Chen:** Validation, Resources, Writing – Review & Editing, Supervision. **Richard A.I. Bethlehem:** Validation, Resources, Writing – Review & Editing. **Jakob Seidlitz:** Validation, Resources, Writing – Review & Editing. **Hongzhe Li:** Validation, Formal Analysis, Writing – Review & Editing, Supervision. **Aaron Alexander-Bloch:** Validation, Resources, Writing – Review & Editing, Supervision. **Yong Chen:** Conceptualization, Methodology, Validation, Resources, Investigation, Writing – Review & Editing, Supervision. **Russell T. Shinohara:** Conceptualization, Methodology, Validation, Formal Analysis, Resources, Investigation, Writing – Review & Editing, Supervision, Project administration, Funding acquisition. **Funding:** FH was supported by NIH Medical Scientist Training Program T32 GM07170. Funding sources were not involved in study design, data analysis, manuscript preparation, or submission decisions.

## Abstract

There is growing interest in estimating population reference ranges across age and sex to better identify atypical clinically-relevant measurements throughout the lifespan. For this task, the World Health Organization recommends using Generalized Additive Models for Location, Scale, and Shape (GAMLSS) which can model non-linear growth trajectories under complex distributions that address the heterogeneity in human populations.

Fitting GAMLSS models requires large, generalizable sample sizes, especially for accurate estimation of extreme quantiles, but obtaining such multi-site data can be challenging due to privacy concerns and practical considerations. In settings where patient data cannot be shared, privacy-preserving distributed algorithms for federated learning can be used, but no such algorithm exists for GAMLSS.

We propose distributed GAMLSS (dGAMLSS), a distributed algorithm which can fit GAMLSS models across multiple sites without sharing patient-level data. We demonstrate the effectiveness of dGAMLSS in constructing population reference charts across clinical, genomics, and neuroimaging settings.

## 1 Introduction

Population reference ranges have been widely used as a cornerstone in clinical medicine to quickly screen a large number of subject-specific measurements, including anthropometrics and laboratory assays, in order to identify atypical measurements that may warrant further clinical follow-up.^1–3^ In recent decades, there has been growing interest in modeling such reference ranges, conceptualized as population reference charts. This idea is inspired by pediatric growth charts which highlight that different reference ranges may be applicable to individuals at different stages of life.^4–7^

In order to estimate such population reference charts, also called normative charts, the World Health Organization has recommended the use of Generalized Additive Models for Location, Scale, and Shape (GAMLSS) as a robust and flexible framework for modeling non-linear growth trajectories.^8–11^ GAMLSS extends generalized linear models (GLM) and generalized additive models (GAM) to allow outcome variables to follow a broad family of distributions, including skewed and heavy-tailed distributions as well as zero-inflated distributions.^12^ GAMLSS also supports semi-parametric modeling of not only the mean of the outcome variable, but also the variance, skewness, and kurtosis of the outcome for any given explanatory variables. This capability allows for the prediction of individualized reference distributions and is essential in the context of reference charts, since sources of population-wide heterogeneity – domain shifts, intrinsic population differences, sampling mechanisms, and more – suggest that higher-order moments may vary with age, sex, and other demographic factors. In addition, this added flexibility requires fewer data assumptions and is thus more generalizable to novel phenotypes. Importantly, while GAMLSS models are highly flexible, the semi-parametric nature of GAMLSS-based population reference charts still allows for a high degree of model transparency, interpretability, and inference, features that are advantageous to the adoption of such reference charts in clinical settings.

Though GAMLSS models offer significant advantages in estimating population reference charts, large sample sizes may be required for fitting. This is especially true as the complexity of the model grows in terms of explanatory variables included, increased non-linearity of the smooth terms, and estimation of higher-order moments. In the context of reference charts, even larger sample sizes are necessary to accurately estimate the extreme quantiles that are essential for identifying atypical measurements, such as those below the 2.5 percentile or above the 97.5 percentile.

Ideally, if individual patient data could be easily shared across multiple institutions, reference charts fit on such data could be confidently applied across a large, representative sample from the general population. However, the ability to construct such charts are often limited by privacy and practical challenges, including governmental regulations such as the Health Insurance Portability and Accountability Act (HIPAA) or the General Data Protection Regulation (GDPR), institutional policies on sharing data across institutions, patient consent protocols for research studies, or even collaborations where investigators are interested maintaining full control of their data. In response to the need for managing large-scale, multi-site data, there has been a rise in the establishment of distributed research networks across healthcare systems, where each healthcare system maintains control of their own data, but cooperates within the network to answer broad-ranging questions via distributed learning. Key examples include the Observational Health Data Sciences and Informatics (OHDSI), the National Patient-Centered Clinical Research Network (PCORnet), the Sentinel Initiative, and the NIH-funded Health Care Systems Research Collaboratory.^13–16^

In such distributed research network settings as well as smaller-scale multi-site collaborations where sharing participant-level data is impractical, it is crucial to develop privacy-preserving federated and distributed learning algorithms that only require summary-level statistics or aggregated data to fit the desired model in a distributed manner. While distributed algorithms have been proposed for various models, including generalized linear models and generalized additive models, no such algorithm has been proposed for fitting GAMLSS.^17–22^

To extend population reference charts to include more representative patient populations and provide more accurate reference quantile estimation for clinical biomarkers, we propose distributed GAMLSS (dGAMLSS). dGAMLSS allows for exact distributed fitting of fully-parametric and semi-parametric models for all GAMLSS family distributions. We demonstrate the applicability of dGAMLSS for building population reference charts in clinical, genomics, and neuroimaging settings for outcomes drawn from various GAMLSS family distributions.^4,23,24^ As big data from electronic health records, clinical genomics, and quantitative neuroimaging become more available, representative population reference charts based on such data may enable the development of quantitative screening biomarkers for health and disease.

## 2 Results

Distributed GAMLSS (dGAMLSS) provides machinery for estimating semi-parametric coeffecients across a broad family of GAMLSS distributions defined by up to four parameters, including mean, variance, skewness, and kurtosis, across multiple sites without sharing any patient-level data. Once coefficients are estimated, dGAMLSS allows for inference and estimation of model-based centiles, defined as quantiles multiplied by 100. Notation for dGAMLSS is defined in Table 1.

**Table 1:**
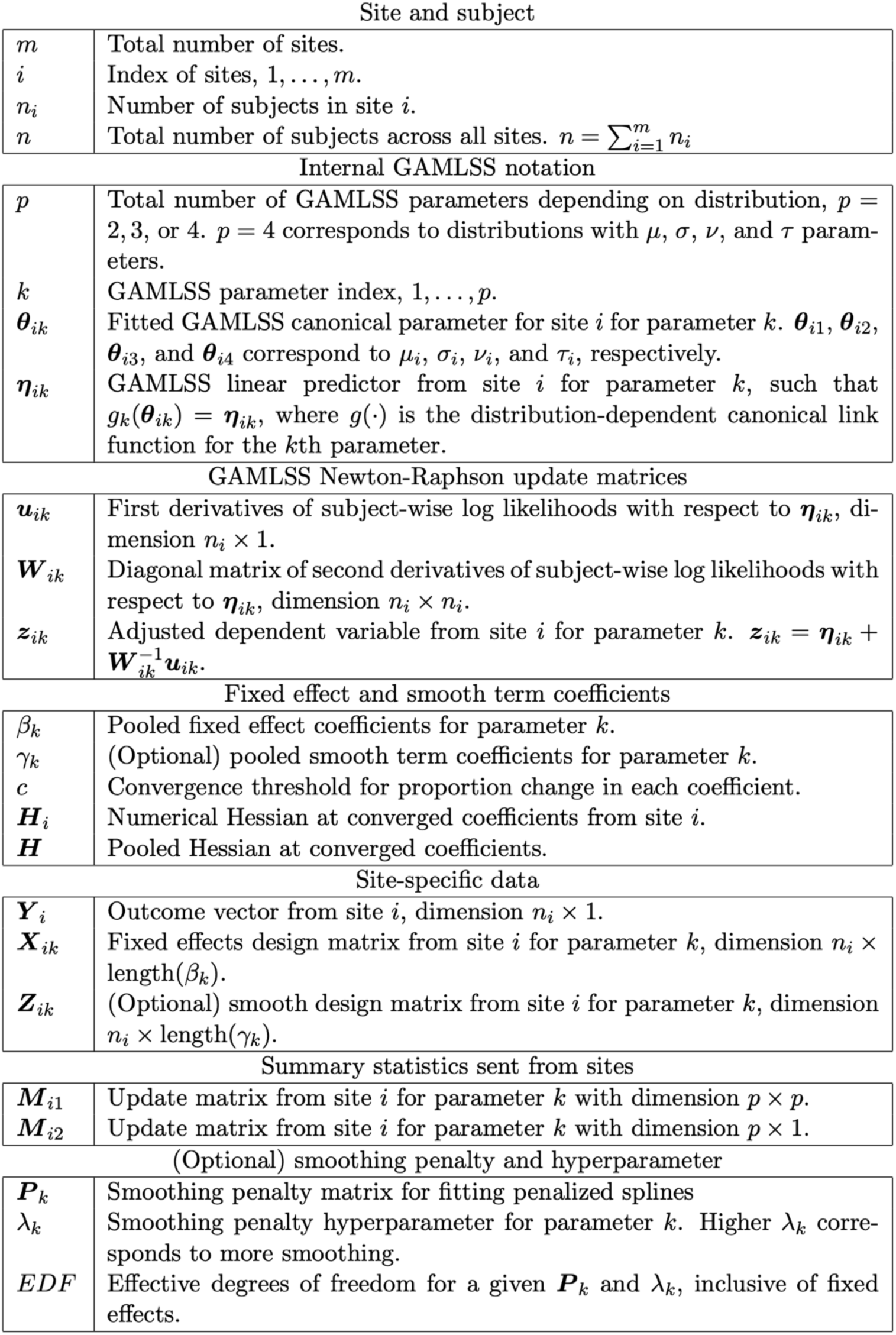
Relevant Notation.

Briefly, the proposed dGAMLSS algorithm adapts the pooled GAMLSS fitting algorithm,^9,10^ which consists of two nested cycles which result in maximization of the likelihood. The outer cycle iterates across the distribution parameters, while the inner cycle updates each parameter’s coefficients using the Newton-Raphson method while keeping coefficients for other parameters constant. Noting that pooled Newton-Raphson updates can be exactly reproduced as the sum of corresponding site-level matrices and substituting as appropriate, dGAMLSS achieves the desired result of privacy-preserving model fitting (Figure 1). Further details are provided in the Methods section and under Algorithms 1 and 2.

**Figure 1:**
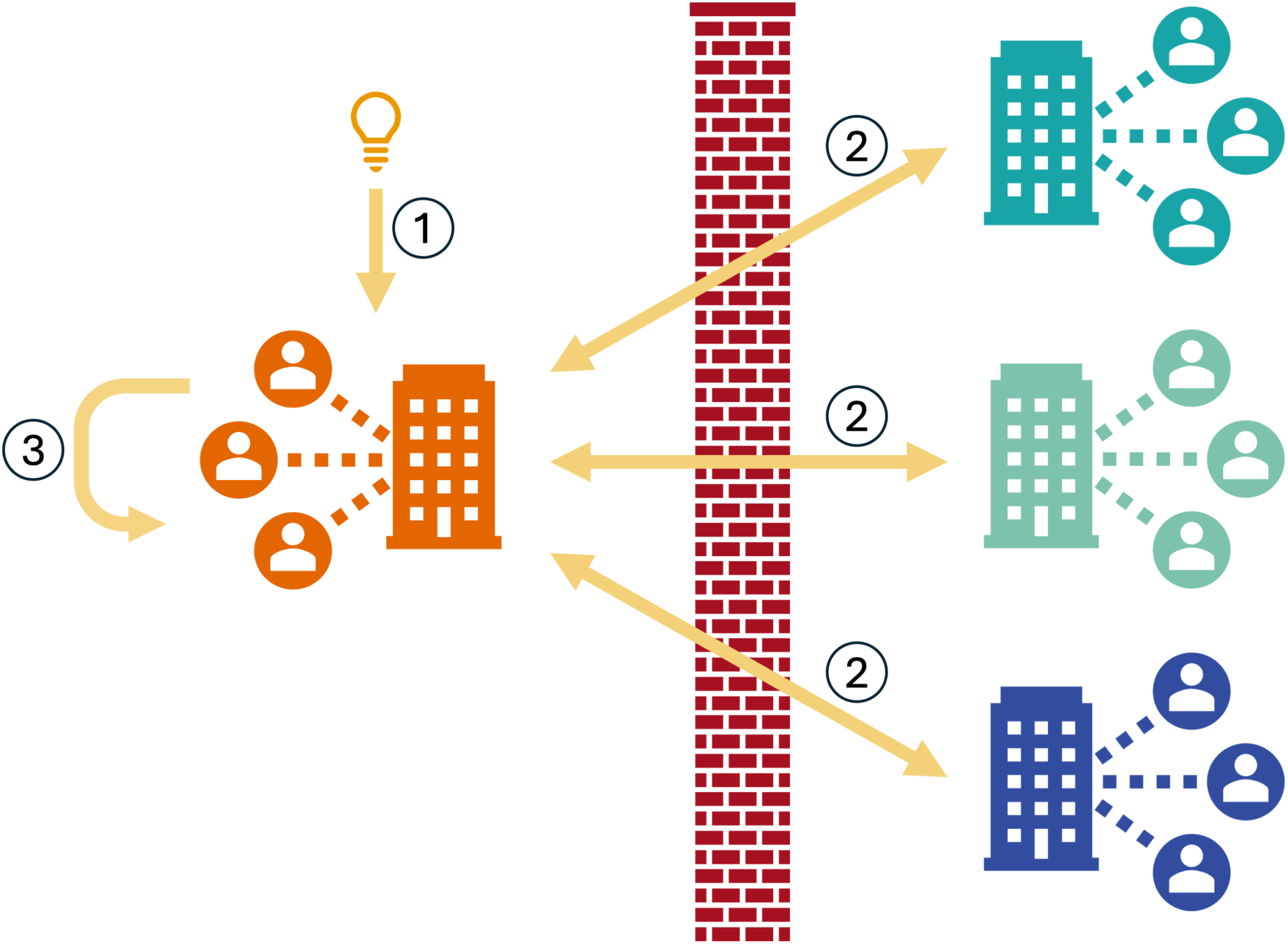
Overview of the dGAMLSS algorithm which enables fitting of GAMLSS models via privacy-preserving federated learning. Fitting occurs via an outer cycle over the parameters and a nested inner cycle of Newton-Raphson updates for parameter-specific coefficients. Broadly, the algorithm is as follows. Step 1: Initialize all coefficients for all parameters and set the outer loop to the *μ* parameter. Step 2: For the given parameter, send current coefficients and receive site-level matrices. Step 3: Calculate coefficient updates for the given parameter. If the updated coefficients are identical to the previous coefficients, advance to the next parameter. Step 4: Repeat Steps 2 and 3 until all coefficients converge for all parameters.

We apply dGAMLSS to estimate body mass index (BMI) charts using electronic health record (EHR) data from the Medical Information Mart for Intensive Care (MIMIC)-IV, microbiome relative abundance charts using data from the Curated Gut Microbiome Metabolome Data Resource, and brain charts using structural magnetic resonance imaging (MRI) data from the Lifespan Brain Chart Consortium (LBCC).^4,23,24^ To demonstrate the various smooth term implementations in dGAMLSS, we use fixed effect smooth terms with regular interval knots in the BMI setting, fixed penalty smooth terms in the microbiome setting, and automated penalty smooth terms in the brain chart setting.

We first provide the fixed effect BMI example to demonstrate the accuracy of dGAMLSS compared to the pooled analysis under a model specification that is relatively straightforward to fit, but may be limited in its ability to fit high amounts of wiggliness between spline knots. We include the fixed penalty microbiome example to show how this limitation can be overcome through the addition of more knots and a penalty, potentially at the cost of a greater number of communication rounds required. The fixed effect and fixed penalty models, though reliant on an assumption of known degrees of freedom that may be impractical in real dGAMLSS applications, are used to illustrate the capacity of dGAMLSS for near-exact fitting and inference compared to identical pooled GAMLSS models. These algorithms may also be useful in scenarios where spline degrees of freedom are roughly known, such as when updating outdated or single-site GAMLSS-based growth charts.

Finally, we provide the automated penalty brain chart example to illustrate how a gold-standard model can be fit using automated penalty selection, with the caveat that this model introduces inferential challenges, requires sites to send additional information per communication round, and may necessitate additional communication rounds.

### 2.1 BMI modeling

The fixed effect dGAMLSS model was fit on an EHR dataset of patients admitted to various intensive care units (ICUs) at Beth Israel Deaconess Medical Center. This dataset included 25112 unique patients from ages 18 to 100 years, where 60.2% of the patients were male and 39.8% of the patients were female. These patients were admitted to nine unique ICUs which, for this example, were treated as unique sites such that patient-level data could not be communicated across ICUs.

We fit a fixed effect smooth model for BMI using the four-parameter Box-Cox power exponential (BCPE) distribution, a generalization of the t-distribution which allows for additional parametric modeling of skewness and kurtosis. The BCPE distribution was chosen based on prior BMI reference charts fit by Rigby and Stasinopoulos and the World Health Organization^25,26^. Each of the four parameters is modeled using a fixed effect smooth term for age at admission and fixed effect terms for sex and admission type.

This model took one communication round to establish the spline basis, 69 communication rounds to converge, and one communication round to obtain inference. The final Bayesian Information Criteria (BIC) of the dGAMLSS model and pooled GAMLSS model were 158575.4 and 158575.3, respectively, indicating effectively equivalent fit.

The dGAMLSS model provided identical inference to the pooled model, with the exception of small numerical differences (Figure 2). Large standard errors for extreme spline bases were observed for both dGAMLSS and pooled GAMLSS models due to a relative lack of observations with non-zero values for those spline bases. dGAMLSS reference BMI charts appeared qualitatively equivalent to pooled reference charts, and both reference charts appropriately reflected BMI trends over the lifespan – larger mean values and variances were observed from ages 25 to 80 while younger and older patients were observed to have comparatively lower BMIs at the median and upper centiles (Figure 3, left). In these age ranges, skewness towards higher BMIs is also observed as well as kurtosis of the upper tails, reflecting the necessity of using a four-parameter distribution such as BCPE for reference BMI chart fitting. Finally, predicted quantiles between dGAMLSS and pooled GAMLSS fits were nearly identical, with the exception of numerical differences (Figure 3, right). The correlation between dGAMLSS and pooled quantile predictions was greater than 0.9999. Smooth patterns observed in the scatterplot are due to numerical differences in spline coefficients. Overall, this analysis demonstrates that, given identical models and spline bases, dGAMLSS provides an exact method for federated learning and inference in the GAMLSS setting.

**Figure 2:**
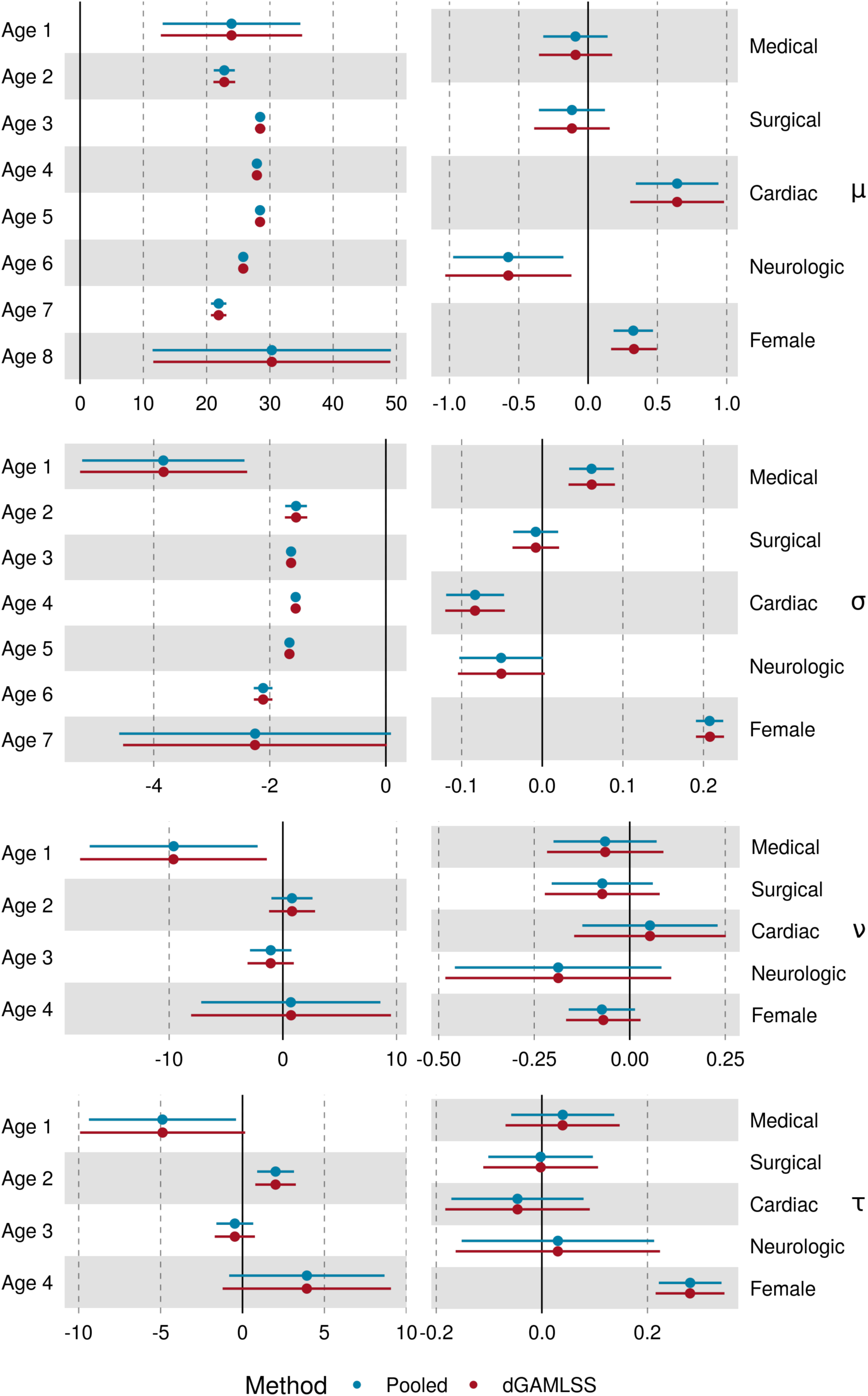
Forest plot of dGAMLSS and pooled GAMLSS coefficients for each parameter. Coefficients for each age spline basis, as well as for sex and admission type, are shown. Age spline bases are numbered from young age to old age. Dots represent point estimates and solid lines represent 95% confidence interval. Vertical dashed lines indicate no effect.

**Figure 3:**
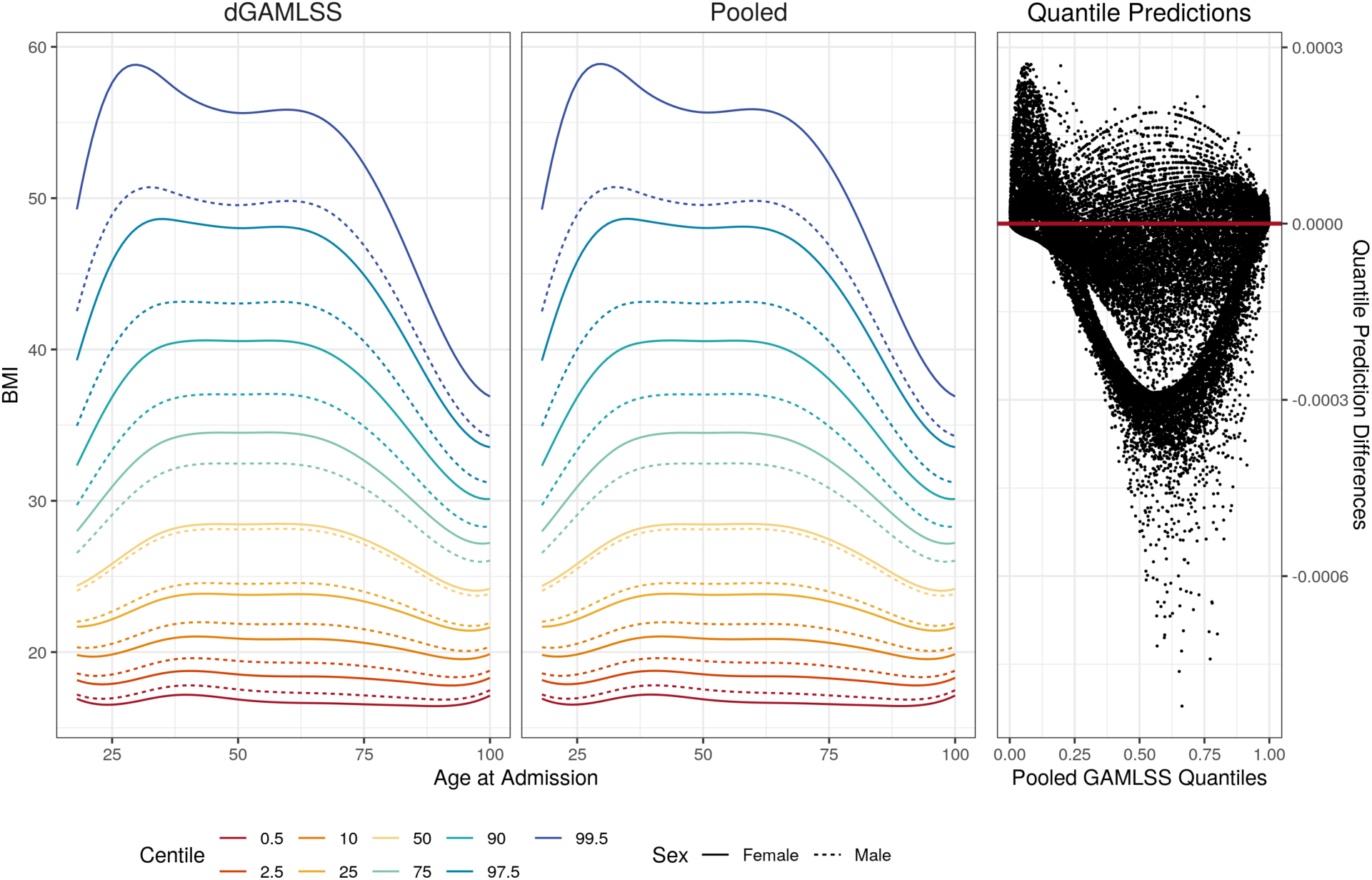
Left: dGAMLSS and pooled GAMLSS reference charts for BMI in ICU patients. Colors represent different centiles, with the orange and purple lines demonstrating a potential reference range defined by the inner 95% of patients. Solid lines represent reference centiles for females while dotted lines represent reference centiles for males. Right: Scatterplot comparison predicted quantiles for every patient. Pooled GAMLSS quantile predictions are shown on the x-axis and differences between dGAMLSS and pooled quantile predictions are shown on the y-axis.

### 2.2 Proteobacteria relative abundance modeling

The fixed penalty microbiome dGAMLSS model used a dataset composed of eleven studies, which were treated as unique sites such that participant-level data could not be transferred between studies. Overall, the dataset included 569 participants from ages 60 days to 83 years where 54.7% were male and 45.3% were female.

We fit a fixed penalty smooth model for Proteobacteria relative abundance using the three-parameter zero-inflated beta distribution, which has been previously proposed for modeling bacteria phyla relative abundances.^27^ The zero-inflated beta distribution is a mixed distribution comprised of the beta and Bernoulli distribution, where the third parameter models the probability of observing zeros. Use of this distribution is supported in our dataset since a meaningful proportion of participants (2.66%) have zero observed Proteobacteria reads.

This model took two communication rounds to establish the spline basis, 117 communication rounds to converge, and one communication round to obtain centiles. The final BIC of the dGAMLSS model and pooled model were highly similar, at -2360.2 and -2365.2, respectively, suggesting minor numerical differences. The estimated degrees of freedom (EDF) of smooth terms for both models were 4.750 for the *μ* term, 4.168 for the *σ* term, and 3.368 for the *ν* term.

dGAMLSS reference Proteobacteria relative abundance charts appeared qualitatively similar to pooled reference charts (Figure 4, left). In these charts, from ages 0 to 40, we observe that around half of all individuals have minimal Proteobacteria relative abundances. However, significant upward skew is observed, as the upper centiles seem to have significantly higher Proteobacteria presence in their microbiomes. For ages above 40, this trend is even more evident, with higher Proteobacteria relative abundances being observed in the upper centiles, while centiles at and below the median remained relatively similar to before. Across ages, females tended to have slightly higher Proteobacteria proportions than males, suggesting more variation of Proteobacteria abundance in females may be normal.

**Figure 4:**
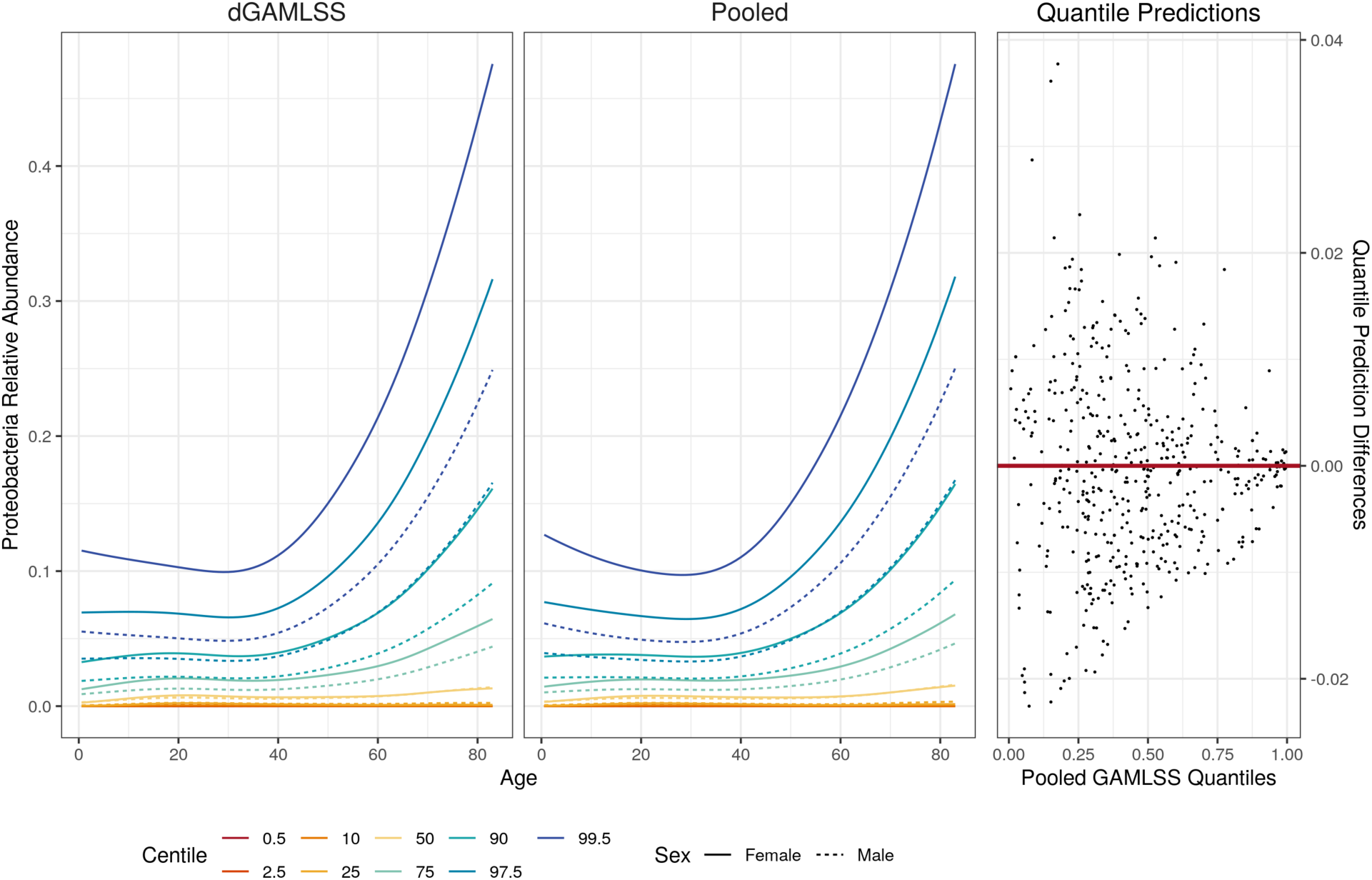
Left: dGAMLSS and pooled GAMLSS reference charts for Proteobacteria relative abundance. Colors represent different centiles, with the orange and purple lines demonstrating a potential reference range defined by the inner 95% of participants. Solid lines represent reference centiles for females while dotted lines represent reference centiles for males. Right: Scatterplot comparison predicted quantiles for every patient. Pooled GAMLSS quantile predictions are shown on the x-axis and differences between dGAMLSS and pooled quantile predictions are shown on the y-axis.

Finally, quantile predictions between fixed penalty dGAMLSS and pooled GAMLSS fits were nearly identical (Figure 5, right) – most differences between predicted quantiles were less than 0.02. The correlation between dGAMLSS and pooled quantile predictions was 0.9994. Overall, this analysis demonstrates that fixed penalty dGAMLSS achieved effectively identical fit when compared to the pooled model.

**Figure 5:**
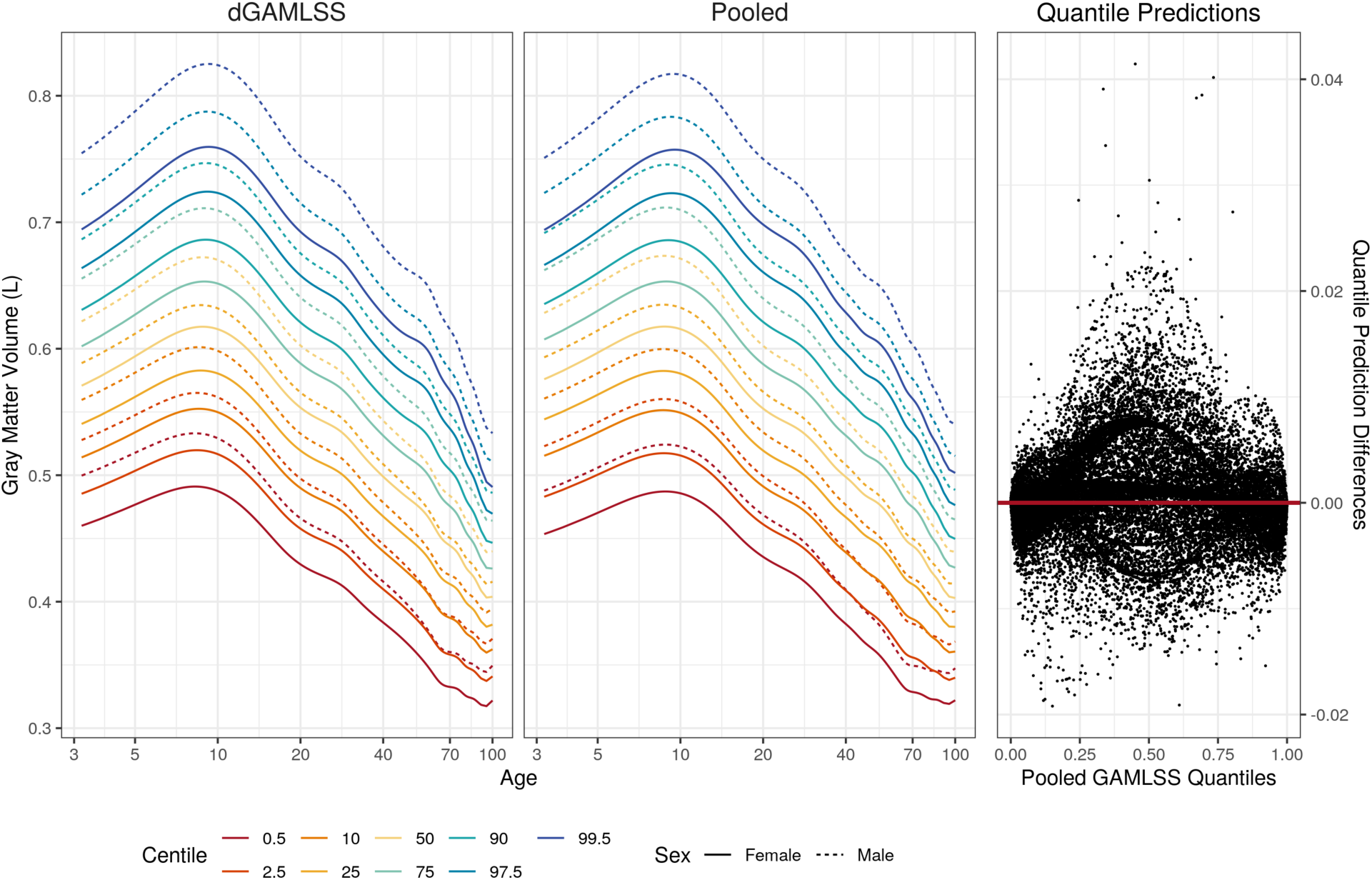
Left: dGAMLSS and pooled GAMLSS reference charts for gray matter volume, in liters, in healthy participants, plotted on a log age scale to better demonstrate large changes in young brains. Colors represent different centiles, with the orange and purple lines demonstrating a potential reference range defined by the inner 95% of participants. Solid lines represent reference centiles for females while dotted lines represent reference centiles for males. Bottom: Scatterplot comparison predicted quantiles for every participant. Pooled GAMLSS quantile predictions are shown on the x-axis and differences between dGAMLSS and pooled quantile predictions are shown on the y-axis.

### 2.3 Gray matter volume modeling

The automated penalty dGAMLSS model was fit on a neuroimaging dataset of 26480 participants from ages 3.2 to 100 years where 48.7% were male and 52.3% were female. Participants spanned 50 studies which were defined as unique sites such that participant-level data could not be transferred from any study.

We fit an automated penalty smooth model for GMV using the three-parameter generalized gamma distribution, chosen based on a prior gold-standard analysis.^4^ The generalized gamma distribution is a generalized version of the gamma distribution allowing for one additional shape parameter. We model the mean and variance parameters with a penalized smooth term for age as well as fixed effects for sex and study. We model the skewness parameter with a penalized smooth term for age and a fixed effect for sex. The B-spline design matrix is specified to have 20 knots, allowing for a high maximum amount of flexibility, such that automated penalization will regularize the effective degrees of freedom. Based on recommendations from Rigby and Stasinopoulos, we use BIC for penalty selection.^10^

The automated penalty dGAMLSS model took one communication round to establish the spline basis and 57 communication rounds to converge. The final automated penalty dGAMLSS model chose 7.32 EDF for the *μ* term, 6.05 EDF for the *σ* term, and 3.01 EDF for the *ν* term. Meanwhile, the pooled GAMLSS model chose for 16.41 EDF for the *μ* term, 5.62 EDF for the *σ* term, and 6.03 EDF for the *ν* term. The final BIC of the dGAMLSS and pooled models were 646260.3 and 646345.7, respectively, indicating the lower EDF selected by the dGAMLSS model may provide better fit than the higher EDF selected by the pooled GAMLSS model. Outside of numerical differences in optimization, additional differences in EDF selection may be due to different knot placement for the spline basis or use of different penalty selection criterion.

dGAMLSS reference GMV charts appeared qualitatively similar to pooled reference GMV charts (Figure 5, left). In these charts, we note a large increase in GMV from around age 3 to 10, steady decreases in GMV from age 10 to 50, and relatively steeper GMV decreases at older ages. Variances seem to be higher at younger ages, and slight skewness towards higher volume is observed across all ages. Male brains tended to have higher GMV. Small differences between dGAMLSS and pooled GAMLSS predictions can be observed for the extreme centiles.

Finally, quantile predictions between automated penalty dGAMLSS and pooled GAMLSS fits were again nearly identical (Figure 5, right) – most differences between predicted quantiles were less than 0.02, though a few subjects demonstrated relatively larger differences. The correlation between dGAMLSS and pooled quantile predictions was 0.9999. Overall, this analysis demonstrates the capacity of automated penalty dGAMLSS to successfully replicate a pooled analysis without *a priori* knowledge of parameter-wise effective degrees of freedom for smooth terms.

## 3 Discussion

In this manuscript, we develop dGAMLSS, an exact, privacy-preserving algorithm for fitting semi-parametric GAMLSS in the federated setting. We show dGAMLSS fits nearly-identical models to pooled GAMLSS in the fixed effect smooth term setting, including coefficient values and Wald-type inference. In the fixed penalty setting, we show dGAMLSS can also achieve similar performance while allowing for potentially higher amounts of non-linearity in certain local regions compared to others. Finally, in the automated penalty setting, we provide machinery for automated penalty selection based on distributed GAIC and GCV criteria and again demonstrate that dGAMLSS is able to reproduce the gold-standard GAMLSS population reference charts.

Further work is necessary to extend dGAMLSS to longitudinal settings by incorporating subject-wise random effects as well as to survival analysis settings by incorporating censored data capabilities.^28–31^ Cross-sectionally, investigation into how to incorporate modern statistical ideas for testing the overall significance of smooth terms in the automated penalty GAMLSS setting would also be beneficial to allow for statistically-sound inference.^32–35^ Notably, this signifance testing limitation also exists in the pooled GAMLSS setting. In a similar vein, the pooled GAMLSS algorithm^10^ utilizes backfitting cycles within the inner Newton-Raphson cycles. This allows GAMLSS to uniquely fit multiple smooth terms for each parameter. However, in the distributed setting, true backfitting is prohibitive – in the setting of just one smooth term, true backfitting requires at least two additional communication rounds per inner cycle. The minimum number of communication rounds per inner cycle increases by two for each additional smooth term, and these communication rounds are multiplied if multiple iterations of backfitting are required in a given inner cycle. While fitting one smooth term per parameter may be adequate in the development of reference charts where age is often the primary smooth term, development of dGAMLSS extensions which allow for communication-efficient backfitting and therefore multiple smooth terms per parameter or interaction smooth terms may be of interest.^36^

Exploration into how dGAMLSS can be made more communication-efficient would allow for further uptake and utility of the algorithm.^20,37,38^ Empirically, most models seem to able to converge in under 100 communication rounds – while fitting such a model is feasible in practice, minimizing communication costs would reduce barriers to use. Other approaches to reducing barriers to dGAMLSS use may be the further development of software specialized to the task of automating communication rounds in federated analysis settings.^39,40^

Privacy-wise, proposed dGAMLSS algorithm follows similar distribution ideas as other federate learning methods, which involves sending two coefficient-based matrices related to the first and second derivatives of overall site-wise likelihoods.^19,21,22^ However, since the dGAMLSS algorithm is not shown to be differentially private, iterative exchanges of summary statistics could lead to incremental information disclosure and increase re-identification risk.^41^ There may be increased privacy risk for sites with small sample sizes relative to the number of covariates, individuals with extreme values for the outcome or one or more covariates, or individuals who have health information in outside databases and could be identified via linking to these outside databases. Such risks are also elevated in the automated penalty setting, since each round of penalty selection requires communication of the above matrices for each set of coefficient locations across a grid of proposed penalties.

Ultimately, we demonstrate the utility of dGAMLSS for building population reference charts in clinical, genomics, and neuroimaging settings where outcomes follow drastically different GAMLSS-family distributions. In the context of modern EHRs and big data efforts where large amounts of data are readily available within institution-specific databases, dGAMLSS promises the potential for real-time, heterogeneity-aware, privacy-preserving reference charts.

## 4 Methods

### 4.1 Pooled GAMLSS framework

We first describe the pooled GAMLSS framework. In the pooled GAMLSS setting where all patient-level data can be collected across sites and accessed simultaneously, pooled data across all sites are notated using an underscore.

GAMLSS-family distributions are a generalized set of probability distributions defined by at least two and up to four parameters. The first two parameters pertain to the mean and variance, while the third and fourth parameters describe skewness and kurtosis, respectively. Such distributions can be continuous, discrete, or mixed distributions such as zero-inflated distributions, allowing for likelihood-based modeling of a wide range of outcome variables.

In GAMLSS, each *n* × 1 vector of subject-specific distribution parameters, ***θ***__1_, … , ***θ***__*p*_, are modeled by additive terms, potentially under different monotonic link functions, *g_k_*(⋅), and differing sets of covariates.^12^ Thus, for each distribution parameter *k* = 1, … , *p*, the semi-parametric parameter-specific GAMLSS model is defined as the following GAM:

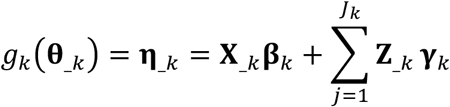

and the goal is to estimate each of the ***β**_k_* and ***γ**_k_* in order to maximize the overall likelihood of observing all subjects’ outcomes given their subject-specific predicted distributions. This model generalizes the GLM to distributions outside of the exponential family and generalizes the GAM such that all parameters, including mean, variance, skewness, and kurtosis, can be modeled in terms of both fixed and smooth terms.

The Rigby and Stasinopoulos (RS) algorithm consists of two cycles which result in maximization of the pooled likelihood with respect to the fixed effect and smoothing coefficients. The outer cycle iterates across the *p* GAMLSS distribution family parameters. Meanwhile, the inner cycle performs Newton-Raphson updates on the outer-cycle parameter-wise coefficients while keeping coefficients for all other parameters constant. In a pooled setting where all data are available, each multivariate Newton-Raphson update can be analytically solved using weighted least squares on the adjusted dependent vector ***z***__*k*_ with diagonal weight matrix ***W***__*k*_, where the underscore represents that the matrices are pooled across all *m* sites. The pooled RS algorithm is described in Appendix B and Appendix C by Rigby and Stasinopoulos in 2005.^10^

### 4.2 Distributed GAMLSS via the RS algorithm

The distributed RS algorithm adapts the pooled RS algorithm, noting that ***z***__*k*_ and the diagonal of ***W***__*k*_ are easily horizontally partitioned into site-specific ***z****_ik_* and ***W****_ik_*. Thus, updates can easily be distributed across sites – as long as each site *i* sends matrices *M_i_*_1_ and *M_i_*_2_ to a central site, the exact pooled weighted least squares solution can be obtained with one round of communication per inner cycle iteration (Table 1). Thus, dGAMLSS is an exact solution for federated learning of GAMLSS models.

Each inner cycle is complete when all parameter-wise coefficients have converged when compared to the prior Newton-Raphson update. The outer cycle is complete when all coefficients across all parameters have converged when compared to their values at the end of the previous outer cycle. Though pooled GAMLSS defines convergence via global deviance change, dGAMLSS defines convergence using coefficient values since global deviance estimation is delayed by one communication round. A given coefficient is considered to be converged when the proportion change between its current and previous value is smaller than some value, *c*. Larger values of *c* allow for faster, but less stable, convergence while smaller values of *c* result in stable convergence, but require more iterations. In pooled GAMLSS, *c* can be made arbitrarily small since additional iterations are processed internally on a central server, which is computationally cheap compared to transferring summary statistics across sites. In the proposed dGAMLSS algorithm, we set *c* = 0.05 to be well-balanced with respect to stability and communication cost. The exact distributed RS algorithm is provided in Algorithms 1 and 2.

#### Algorithm 1 Distributed RS algorithm, fixed degrees of freedom

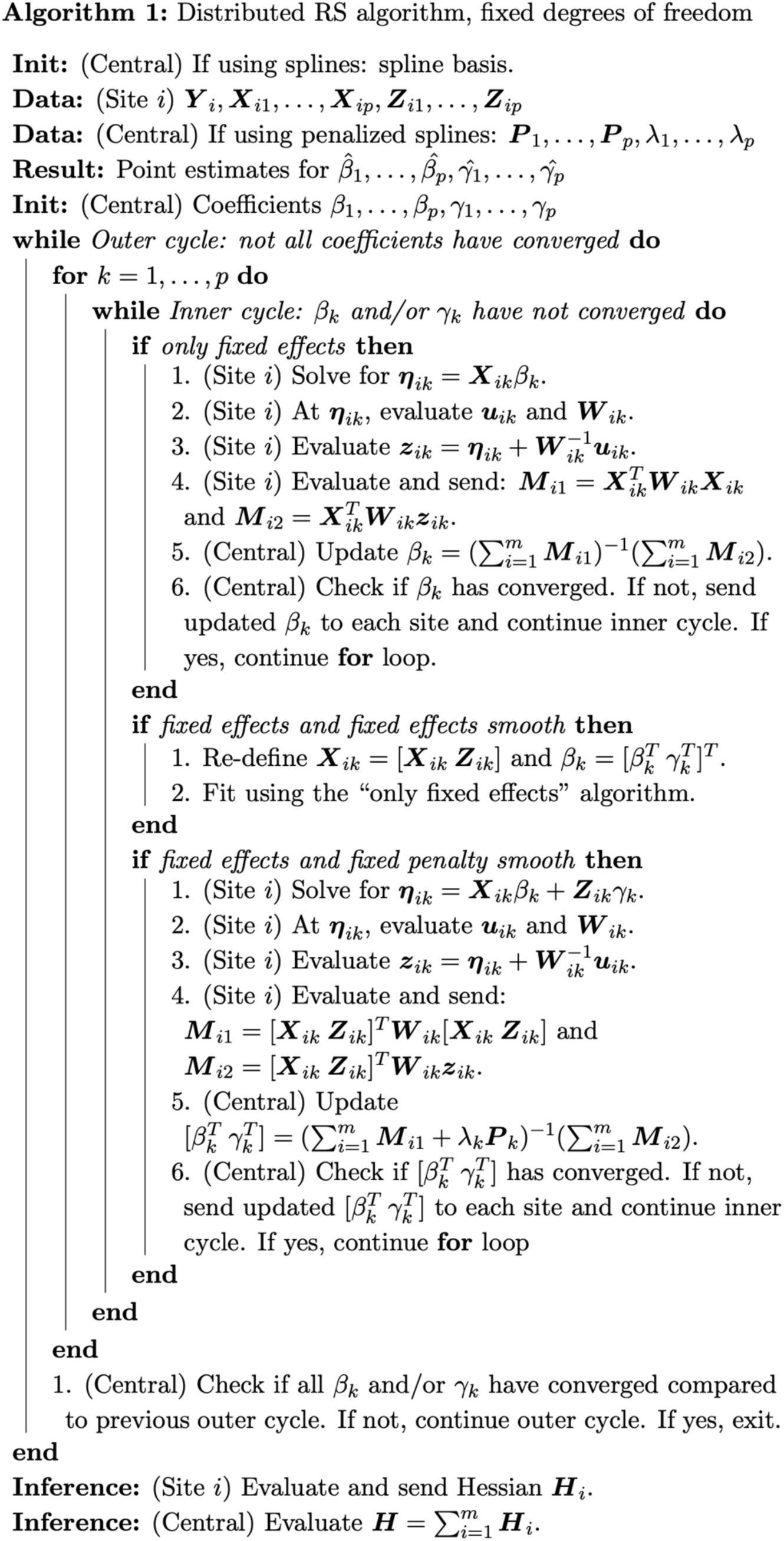

#### Algorithm 2 Distributed RS algorithm, automated penalty selection

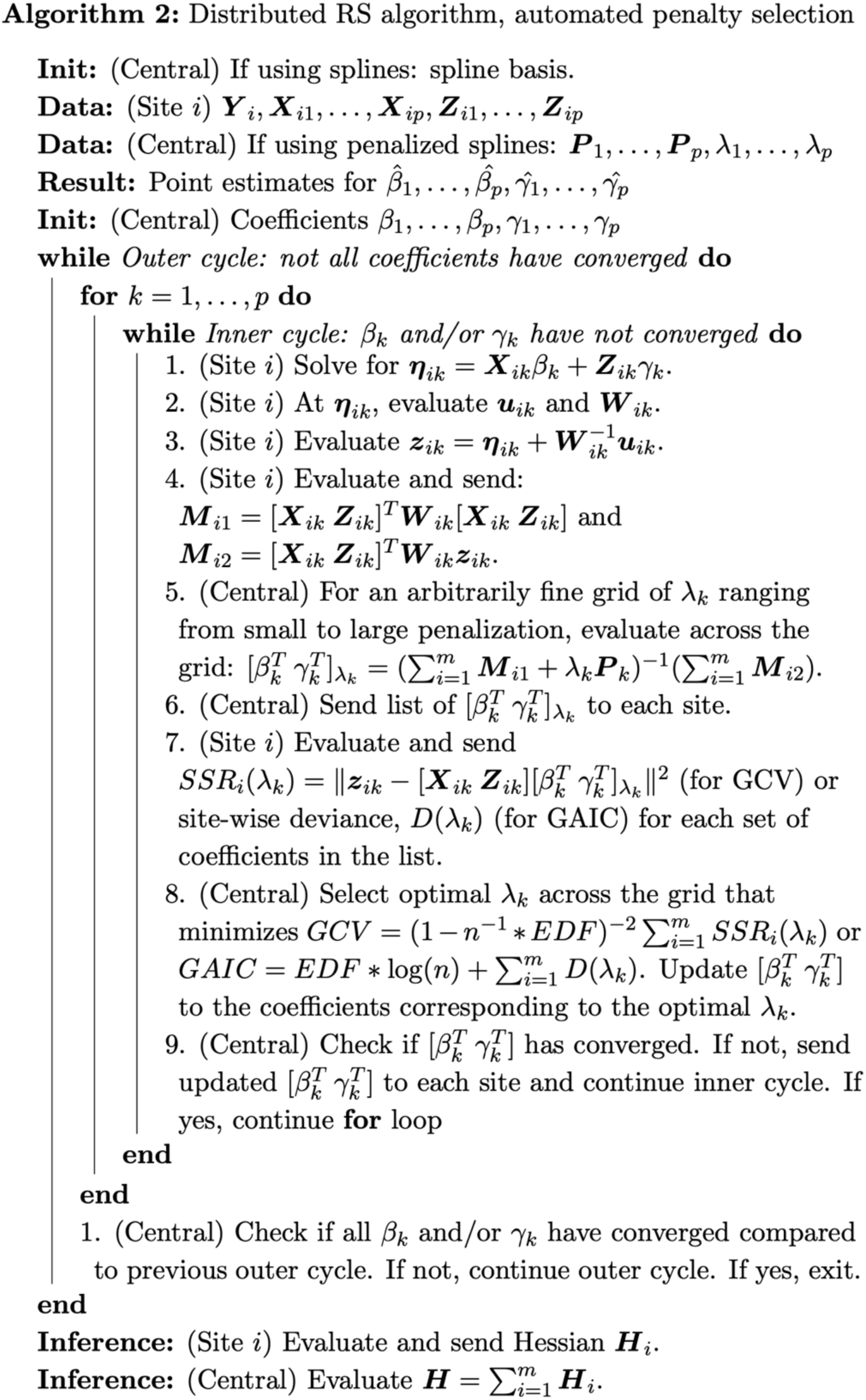

### 4.3 Distributed smooth terms

We provide three distributed RS approaches of increasing complexity for fitting up to one smooth term per parameter. We first discuss using fixed effect smooth terms, where wiggliness of the smooth term is directly controlled by limiting the number of spline knots. Next, we describe fixed penalty smooth terms, where the spline basis is specified with more knots than necessary and wiggliness of the smooth term is controlled via a known penalty hyperparameter. Finally, we propose an algorithm for fully-automated penalty selection for smooth terms using either generalized Akaike Information Criterion (GAIC) or generalized cross-validation (GCV). For all smooth terms in dGAMLSS, we use B-splines, but other types of splines can also be used.

#### 4.3.1 Distributed knot placement and smooth design matrix specification

To fit a given smooth term, dGAMLSS requires that a common spline basis be used across all *i* sites. This basis must be chosen such that the global minimum and global maximum for the relevant covariate are contained within the boundary knots of the spline, an identical number of knots are used across sites, and knots are placed at identical locations across sites. Knots can either be placed at regular intervals over the range of the covariate values or at approximate quantiles of the relevant covariate for maximum efficiency. If fixed effect smooth terms are used, either knot placement approach can be used – regular interval knots are easier to place, while approximate quantile are more efficient but require additional information to be sent in the first communication round. For fixed effect smooth terms, a total of *EDF* − 2 knots should be placed, including the two boundary knots, in order to achieve a given desired EDF. If fixed penalty smooth terms are used, regular interval knot placement is adequate, given that enough knots are placed, and EDF is controlled using a penalty hyperparameter, discussed below.

One round of pre-fitting communication is required to specify the spline basis. If regular interval knots are desired, each site must send site-specific ranges for the relevant covariate such that the global range can be calculated. Knots can then be placed at regular intervals along this global range by the central site, and these knot locations can be sent back to other sites. If approximate quantile knots are desired, relevant summary statistics should also be simultaneously sent in this communication round, such as site sample size as well as site-specific covariate means and standard deviations (if the covariate distribution is thought to be nearly normal) or some number of quantiles (if the covariate distribution is thought to be non-normal). If site-specific covariate means and standard deviations are sent, these summary statistics and the site sample sizes can be used by the central site to estimate the pooled covariate mean and standard deviation and therefore obtain approximate quantiles. If site-specific quantiles, such as deciles are sent, the central site can simulate an appropriate number of uniformly distributed observations for each site within each decile, pool simulated observations across sites, and obtain approximate quantiles. These approximate quantiles can be sent back to other sites as the final knot locations. Once knots are placed, a common spline basis across all sites is achieved.

For fixed penalty or automated penalty smooth terms, additional information is necessary to learn the relationship between *λ_k_* and smooth term EDF, where each site must provide one additional round of pre-fitting communication containing site-specific matrices [***X****_ik_* ***Z****_ik_*]*^T^*[***X****_ik_* ***Z****_ik_*] for each parameter containing penalized smooths. To do so, a common spline basis is defined as above, and a joint penalty matrix across the fixed effects and smooth term is automatically specified based on the spline basis.^42^ This joint penalty matrix for parameter *k*, ***P****_k_*, is a four-block square matrix of size length(***β**_k_*) + length(***γ**_k_*), where the bottom right block matrix is the length(***γ**_k_*) × length(***γ**_k_*) spline penalty matrix and the other three block matrices are ***0*** matrices of appropriate size.

This joint specification of the penalty matrix allows for direct fitting of the fixed effects and smooth terms without backfitting. Note that, if orthogonalization is performed for the smooth design matrix, the standard B-spline penalty matrix must be appropriately transformed using the ***R***^−1^ matrix from above. Finally, the following formula is used to obtain the final EDF of the smooth term for any given penalty hyperparameter *λ_k_*:

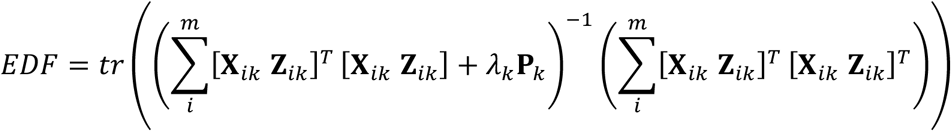

Once necessary smooth design matrices and penalty matrices, as necessary, are defined, model fitting can proceed. Model fitting for fixed effect and fixed penalty models are described in Algorithm 1. Model fitting for automated penalty models is described in Algorithm 2. Notably, for automated penalty models, the default option of maximum likelihood penalty selection cannot be efficiently used in the distributed setting since the maximum likelihood algorithm requires iterative optimization of the penalty hyperparameter. Meanwhile, GAIC and GCV algorithms simply require selection of a penalty hyperparameter such that GAIC or GCV are minimized. In the distributed setting, testing GAIC or GCV across a grid of many hyperparameter values can be performed in one communication round.

### 4.4 Distributed inference

Once the distributed RS algorithm has converged and coefficient point estimates are obtained, estimated GAMLSS quantiles can be easily obtained for the reference population as well as any given new data. For reference population quantiles, the central site can simulate data on a grid across all covariates. These simulated covariates can be combined with coefficient point estimates to obtain predicted distributions for each set of covariates, and reference quantiles can be obtained from these predicted distributions. Covariates from new data can be directly used to estimate predicted distributions, and quantiles for outcomes from new data can be estimated.

To perform inference, one additional round of post-fitting communication is necessary. The coefficient point estimates are sent to each site, and each site returns numerical site-wise likelihood-based Hessians, ***H****_i_*, simultaneously evaluated across all parameters. The global covariance matrix, 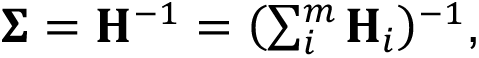 is calculated by inverting the sum of site-specific Hessians. The diagonal of this covariance matrix is used in GAMLSS to obtain standard errors and t-statistics for each coefficient and therefore perform Wald-type inference. In this round of communication, site-specific deviances for the final model are also returned and summed to obtain the global deviance of the model.

For models using fixed effect smooth terms and fixed penalty smooth terms, inference on the overall effect of the spline can be performed via likelihood ratio test comparing the global deviance of a full model with smooth terms to the global deviance of a reduced intercept-only model, where the null distribution of the likelihood ratio test is 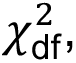 where df is the number of columns of the fixed effect smooth term or the EDF of the fixed penalty smooth term being tested. Note that if multiple parameters contain smooth terms, inference for a given parameter requires a reduced intercept-only model for that parameter only; other parameters must still contain their full smooth terms. To reduce overall communication rounds, if likelihood ratio test inference is desired, reduced models should be fit simultaneously to full models.

For models using automated penalty smooth terms, use of the likelihood ratio test discussed above may be liberal since it does not appropriately account for the data-driven choice of *λ_k_*.^32–35^ Exact inference is not implemented for smooth terms in this context. However, in many GAMLSS settings where the goal is to predict the quantiles of new data given their covariates, optimally-penalized smooth terms may be desirable for their accurate model fit, despite challenges with inference.

### 4.5 dGAMLSS applications

#### 4.5.1 Modeling BMI in intensive care unit patients

Patients from the MIMIC-IV database were used to model the relationship between age, sex, and BMI, controlling for different intensive care unit (ICU) types. The MIMIC-IV database has been previously described.^23^ While the MIMIC-IV database may be sub-optimal for developing reference BMI charts applicable to a non-ICU population, the EHR setting provides a compelling scenario for the demonstration of dGAMLSS.

We filtered the MIMIC-IV dataset such that patients who had extreme heights, weights, or BMIs, where extremeness was defined as less than the 1st percentile or greater than the 99th percentile, were excluded. This accounted for data entry errors, such as incorrect measurement units, accidental extra digits, and more, which would otherwise result in a small subset of patients having physiologically implausible heights, weights, or BMIs. Additionally, hospital stays where patients were transferred between ICUs were removed such that each remaining patient was present in exactly one ICU dataset. Finally, if a patient had more than one hospital stay resulting in longitudinal measurements, BMI and admission age from the first hospital stay was retained and following timepoints were discarded, since dGAMLSS requires cross-sectional data.

This filtering resulted in a total of 25112 unique patients who were admitted at ages from 18 to 100 years. 60.2% of the patients were male while 39.8% of the patients were female. These patients were admitted to nine unique ICUs, with the following breakdown: Medical ICU (MICU; n = 4222), Medical/Surgical ICU (MSICU; n = 3522), Surgical ICU (SICU; n = 2584), Trauma Surgical ICU (TSICU; n = 2616), Coronary Care Unit (CCU; n = 3102), Cardiac Vascular Intensive Care Unit (CVICU; n = 8269), Neuro Stepdown Unit (n = 140), Neuro Intermediate Care Unit (n = 340), Neuro Surgical ICU (NSICU, n = 317). To control for the relationship between admission type and BMI, patients were defined as medical, surgical, neurologic, or cardiac based on which ICU the patients were admitted to. For example, patients admitted to the MSICU were defined as both medical and surgical patients. For the purposes of this demonstration, each ICU was defined as a unique site such that patient-level data could not be communicated across ICUs.

We fit a fixed effect smooth model for BMI using the four-parameter Box-Cox power exponential (BCPE) distribution, a generalization of the t-distribution which allows for additional parametric modeling of skewness and kurtosis. Each of the four parameters is modeled using a fixed effect smooth term for age at admission and fixed effect terms for sex and admission type. For each parameter, non-orthogonalized, regular-interval B-spline design matrices are defined using the following total number of knots: *μ*: 6, *σ*: 5, *ν*: 2, and *τ*: 2. These knots are chosen based on the degrees of freedom given by the gold standard model and are used to demonstrate that, if optimal degrees of freedom are known, dGAMLSS can successfully fit the desired model.

In this analysis, we compare the fitted dGAMLSS model to a pooled GAMLSS model using the same fixed effect spline basis in order to show that dGAMLSS accurately reproduces pooled GAMLSS output when splines are identically defined. We analyze dGAMLSS coefficient and standard error estimates in comparison to those from the pooled GAMLSS models to assess the validity of distributed inference. Additionally, we plot dGAMLSS predicted quantiles against pooled GAMLSS predicted quantiles for each observation to assess accuracy of predicted quantiles. Finally, we construct dGAMLSS reference charts and compare them to pooled GAMLSS reference charts.

#### 4.5.2 Microbiome relative abundance modeling

Microbiome data from the gut microbiome-metabolome dataset collection were used to model how Proteobacteria relative abundance changes across age and sex. Processing of this dataset is previously described.^24^ Of the fourteen datasets included in the collection, two datasets were excluded due to lack of sex covariate and one dataset was excluded due to lack of exact age covariate. Participants outside of the control group were excluded so that reference Proteobacteria relative abundance could be modeled. When more than one microbiome sample was available for a given participant, the earliest sample was retained, and following measurements were discarded.

In the end, our analysis included 569 participants from 60 days to 83 years of age. In this sample, 54.7% were male and 45.3% were female. The eleven studies were treated as unique sites such that participant-level data could not be transferred between studies. The smallest study contained eight subjects while the largest study contained 89 subjects.

We fit a fixed penalty smooth model for Proteobacteria relative abundance using the three-parameter zero-inflated beta distribution, which has been previously proposed for modeling bacteria phyla relative abundances.^27^ The zero-inflated beta distribution is a mixed distribution comprised of the beta and Bernoulli distribution, where the third parameter models the probability of observing zeros. Use of this distribution is supported in our dataset since a small proportion of participants (2.66%) have zero observed Proteobacteria reads.

We model the mean parameters with a fixed penalty smooth term for age as well as fixed effects for sex and dataset. We model the variance and zero-inflation parameters with a fixed penalty smooth term for age and a fixed effect for sex. The B-spline design matrix is specified to have 20 knots, allowing for a high maximum amount of flexibility. Penalties are manually selected such that the overall EDF of each smooth term is equivalent to that of the pooled GAMLSS model. While this approach to choosing penalties is not feasible in a real application of dGAMLSS, we use this approach to demonstrate the capability of dGAMLSS to fit penalized smooths when a gold-standard penalty is known.

We compare the fitted dGAMLSS model to the pooled GAMLSS model fit using penalized B-splines with default maximum likelihood penalty selection. We plot dGAMLSS predicted quantiles against pooled GAMLSS predicted quantiles for each observation to assess accuracy of predicted quantiles. We also compare dGAMLSS reference charts with pooled GAMLSS reference charts to assess quality of dGAMLSS microbiome charts.

#### 4.5.3 Brain volumetric modeling

Neuroimaging data from a subset of participants in the LBCC were used to model how gray matter volume (GMV) changes across age and sex. Details on how GMV were obtained are previously described.^4^ Exclusion criteria for this subset involved removal of low-quality scans and non-physiologically plausible brain volumes.^43^ Additionally, LBCC data from the UK Biobank was excluded from this analysis due to data sharing challenges.

Ultimately, 26480 participants were included in this analysis, spanning ages 3.2 to 100 years. Of these participants, 48.7% were male and 52.3% were female. Participants spanned 50 studies which, for the purposes of this demonstration, were defined as unique sites such that participant-level data could not be transferred from any study. Study sample sizes ranged such that the smallest included study only contributed 5 participants while the largest included study contributed 7889 participants.

We fit an automated penalty smooth model for GMV using the three-parameter generalized gamma distribution, chosen based on recommendations from Bethlehem et al.^4^ The generalized gamma distribution is a generalized version of the gamma distribution which allows for one additional shape parameter. We model each of the mean and variance parameters with a penalized smooth term for age as well as fixed effects for sex and study. We model the skewness parameter with only a penalized smooth term for age and a fixed effect for sex. The B-spline design matrix is specified to have 20 knots, allowing for a high maximum amount of flexibility, such that penalization will automatically regularize the effective degrees of freedom. Based on recommendations from Rigby and Stasinopoulos, we use BIC for penalty selection.^10^

We compare the fitted dGAMLSS model to the pooled GAMLSS model fit using penalized B-splines with default maximum likelihood penalty selection. We plot dGAMLSS predicted quantiles against pooled GAMLSS predicted quantiles for each observation to assess accuracy of predicted quantiles. We also compare dGAMLSS reference charts with pooled GAMLSS reference charts to assess quality of dGAMLSS brain charts.

## 4.6 Data and code availability

An R package containing all code for running dGAMLSS, performing distributed inference, and building distributed reference charts is provided at: https://github.com/hufengling/dGAMLSS. All code for analysis and manuscript preparation is provided at: https://github.com/hufengling/dGAMLSS_analyses. Included in the dGAMLSS package and analysis code are vignettes demonstrating how to set up dataframes and R objects for distributed computation as well as functions simulating how coefficient updates are performed as the central site.

The MIMIC-IV database is publically available via PhysioNet.^44,45^ The microbiome dataset is publically available and was accessed via: https://github.com/borenstein-lab/microbiome-metabolome-curated-data.^24^ Availability of the LBCC database is managed at the discretion of each primary study and is previously described.^4,43^ Links to open and semi-open access datasets are listed at: https://github.com/brainchart/Lifespan.

## Notes

**Disclosures and Conflicts of Interest:** RTS receives consulting income from Octave Bioscience and compensation for reviewership duties from the American Medical Association. JS, RAIB and AA-B hold shares in Centile Bioscience, and JS and RAIB are directors of Centile Bioscience.

### Competing Interest Statement

RTS receives consulting income from Octave Bioscience and compensation for reviewership duties from the American Medical Association. JS, RAIB and AA-B hold shares in Centile Bioscience, and JS and RAIB are directors of Centile Bioscience.

